# *Juvenile* AAV-Mediated MEF2C Gene Replacement Ameliorates Selected Phenotypes in *Mef2c*-Haploinsufficient Mice

**DOI:** 10.64898/2026.08.14.744746

**Authors:** Zhuolei Jiao, Chengjie Yu, Tianshu Li, Yuanting Lai, Yifan Yang, Yiting Yuan, Yuefang Zhang, Gongjia Tao, Junwen Wang, Zilong Qiu

## Abstract

MEF2C haploinsufficiency syndrome is a severe neurodevelopmental disorder for which no disease-directed treatment is available. We investigated whether neuron-directed adeno-associated virus (AAV) delivery of a functional MEF2C coding sequence during the juvenile period could modify disease-relevant phenotypes in mice heterozygous for a *Mef2c* exon 4 deletion. Transcript-level analysis identified a brain-enriched MEF2C isoform containing the α1 and β regions (nMEF2C) and a skeletal-muscle-enriched isoform containing α2 but lacking β (mMEF2C). Separate human-synapsin-driven AAV vectors encoding either isoform were administered at postnatal day 28. Control-treated *Mef2c* heterozygous mice retained baseline sociability but lacked social-novelty preference. Mice treated with either nMEF2C or mMEF2C displayed social-novelty preference and improved selected responses to a new social partner, whereas open-field effects were limited. nMEF2C replacement also corrected dark-phase wakefulness and non-rapid eye movement sleep abnormalities and modified selected state-dependent electroencephalographic ratios, without broadly changing absolute band amplitudes or social-contact electroencephalographic activity. Atlas-based whole-brain mapping revealed region-selective reductions in parvalbumin-immunoreactive profiles; direct statistical evidence of cellular rescue was confined to the secondary motor area after nMEF2C treatment. These findings show that selected MEF2C-dependent phenotypes remain modifiable during the juvenile period and support further optimization of MEF2C gene replacement with respect to isoform, dose, expression control, and cellular targeting.

## Introduction

MEF2C-related neurodevelopmental disorder, also termed MEF2C haploinsufficiency syndrome (MCHS), is caused predominantly by de novo heterozygous deletions or pathogenic variants that reduce myocyte enhancer factor 2C (MEF2C) dosage or function. Affected individuals commonly have severe developmental delay and intellectual disability, absent or limited speech, hypotonia, motor impairment, stereotypic movements, autistic features, seizures, electroencephalographic abnormalities, and sleep disturbances.^1–4^ Current management is supportive, and no approved therapy restores functional MEF2C.^4^

MEF2C is an activity-regulated MADS/MEF2-family transcription factor that governs neural progenitor differentiation, neuronal maturation, synapse development, and plasticity.^5,6^ Mouse models implicate excitatory and inhibitory neurons and neuroimmune populations in the composite phenotype of MEF2C deficiency.^6–8^ Reduced parvalbumin (PV) immunoreactivity has been reported in some *Mef2c*-haploinsufficient models,^9^ whereas GABAergic-cell MEF2C hypofunction impairs PV-interneuron physiology.^8^ MEF2C also contributes to cortical transcriptional and electrophysiological responses to sleep loss,^10^ suggesting that postnatal circuit dysfunction may remain therapeutically modifiable.

Postnatal NitroSynapsin treatment improved cellular, electrophysiological, and behavioral abnormalities in *Mef2c*-haploinsufficient mice.^9^ DNA and RNA editing have also ameliorated phenotypes caused by the specific *Mef2c* p.Leu35Pro allele.^11,12^ However, mutation-specific editing is not directly applicable to whole-gene deletions or many other loss-of-function alleles. Delivery of a functional MEF2C coding sequence may therefore offer a complementary, potentially genotype-flexible approach, although the dosage sensitivity and context-dependent activity of MEF2C require careful control of targeting and expression.

Here, we tested neuron-directed AAV-mediated MEF2C gene replacement initiated at postnatal day 28 in mice heterozygous for a *Mef2c* exon 4 deletion. We compared brain-enriched and muscle-enriched human MEF2C isoforms across behavioral and atlas-registered PV-immunoreactivity outcomes and further evaluated sleep–wake and electroencephalographic effects of the brain-enriched isoform.

## Materials and Methods

### Study design

This study evaluated whether neuron-directed adeno-associated virus (AAV)-mediated replacement of two human MEF2C isoforms could ameliorate phenotypes associated with *Mef2c* haploinsufficiency. The brain-enriched isoform was designated nMEF2C, whereas the muscle-enriched isoform was designated mMEF2C. Separate cohorts of *Mef2c* heterozygous mice received AAV expressing nMEF2C, mMEF2C, or EGFP as a vector control. Wild-type (WT) littermates served as reference controls.

Vectors were administered at postnatal day 28 (P28). Behavioral testing began at P56 and included the open-field, three-chamber social-interaction repetitive social-interaction tests, and novel object tests. Following behavioral testing, the nMEF2C cohort underwent 24-hour EEG/EMG recording and sleep–wake analysis. The nMEF2C and mMEF2C cohorts were used for atlas-based whole-brain analysis of parvalbumin (PV) immunoreactivity at P130. Sample sizes for each experiment are reported in the corresponding figure legends.

### Analysis and selection of human MEF2C isoforms

Transcript-level MEF2C expression (ENST00000504921.7 and ENST00000340208.9) was examined using publicly available human tissue transcriptomic data from https://www.gtexportal.org/home/gene/MEF2C/exonExpressionTab. The corresponding human protein sequences were obtained from UniProt. ENST00000504921.7/UniProt Q06413-1, which was enriched in brain tissues, was designated nMEF2C. ENST00000340208.9/UniProt Q06413-5, which was enriched in skeletal muscle, was designated mMEF2C. Protein sequences were aligned using https://www.uniprot.org/align. The two isoforms share the conserved N-terminal MADS and MEF2 DNA-binding domains but differ in their alternatively spliced α regions and in the presence or absence of the β segment. nMEF2C contains the α1 region and β segment, whereas mMEF2C contains the α2 region and lacks the β segment.

### Animals and ethical approval

*Mef2c* heterozygous mice and WT littermates were maintained on a C57/BL6 background in a specific-pathogen-free facility. Mice were housed under a 12-hour light/12-hour dark schedule, with lights on at 7am and off at 7pm. Food and water were provided ad libitum. Mice of both sexes were included, animals were distributed across treatment groups with sex and litter considered during allocation.

All procedures were reviewed and approved by the Institutional Animal Care and Use Committee of Songjiang Hospital Affiliated to Shanghai Jiao Tong University School of Medicine under protocol “ACE-004-2025”.

### Generation and genotyping of Mef2c heterozygous mice

The mutant *Mef2c* allele was generated by deleting exon 4 using CRISPR/Cas9. The deletion boundaries were confirmed by sanger sequencing. Founder animals were backcrossed to C57BL/6 for 2 generations before experimental breeding.

Genotyping PCR was performed with primers flanking the exon 4 deletion: forward, 5′ - AGTCCCCAGTAGTTCCCCAGTG-3′; reverse, 5′-TCTGTGAGAATGGTCTGGGCTCTGGA-3′. PCR products were resolved by agarose-gel electrophoresis. Products of 1,587 and 256 bp identified the WT and deleted alleles, respectively. Heterozygous animals carried both alleles.

### AAV vector construction and production

Human nMEF2C and mMEF2C coding sequences were cloned separately downstream of the human synapsin promoter. Each expression cassette contained, in the 5′-to-3′ direction, an AAV2 inverted terminal repeat, the human synapsin promoter, a Kozak sequence, the nMEF2C or mMEF2C coding sequence, a T2A self-cleaving peptide, EGFP, the human growth hormone polyadenylation signal, and a second AAV2 inverted terminal repeat. The T2A sequence permitted production of MEF2C and EGFP as separate proteins from a single transcript. The control vector expressed EGFP under the same promoter without a MEF2C coding sequence. The nMEF2C and mMEF2C coding sequences corresponded to ENST00000504921.7 and ENST00000340208.9, respectively. Recombinant vectors were packaged in an AAV-PHP.EB capsid by PackGene inc.

### AAV administration

At P28, *Mef2c* heterozygous mice received AAV-hSyn-nMEF2C-T2A-EGFP, AAV-hSyn-mMEF2C-T2A-EGFP, or AAV-hSyn-EGFP by intravenous tail-vein injection, as previously described.^11^ Each animal received 1 × 10^13^ vector genomes in a total volume of 100 μL. Animals were returned to their home cages after recovery and monitored for 1 hour.

### Behavioral testing

Mice were tested beginning at P56 during the light phase. Before testing, animals were handled on three consecutive days and acclimated to the testing room for at least 30 minutes. Assays were separated by 24–48 hours. Apparatuses and stimulus enclosures were cleaned between animals. Behavior was video recorded and quantified using homemade python code by investigators blinded to genotype and treatment.

### Open-field test

Locomotor and center-exploration behavior was assessed in a square arena measuring 40 × 40 cm. The central region was defined as 20 × 20 cm. Each mouse was recorded during a 10-minute test session. Mean velocity, velocity in 1-minute bins, and time spent in the central region were calculated. Illumination at the arena floor was 50 lux, and the mouse was initially placed at the center of the open-field.

### Three-chamber social-interaction test

The apparatus contained two side compartments connected to a central compartment. Mice were habituated to the empty apparatus for 10 minutes 1 day before testing and immediately before testing.

During the social-approach phase, an unfamiliar stimulus mouse was placed beneath a wire enclosure in one side compartment, and an identical empty enclosure was placed in the opposite compartment. The subject mouse was introduced into the center and allowed to explore for 10 minutes. During the subsequent social-novelty phase, a second unfamiliar mouse was placed beneath the previously empty enclosure, whereas the first mouse remained as the familiar stimulus. Exploration was recorded for another 10 minutes. The sides assigned to the social and empty or novel stimuli were counterbalanced.

Interaction time was defined as orientation within 3.5 cm from each enclosure and was quantified using homemade automated tracking python code. Stimulus mice were same sex with the subject, and were habituated to the enclosures at least 1 hour before testing.

### Repetitive social-interaction test

Each subject underwent four 3-minute encounters with the same unfamiliar stimulus mouse. Consecutive encounters were separated by 5-minute intervals. In a fifth 3-minute trial, the familiar stimulus mouse was replaced with a new unfamiliar mouse. Testing was conducted in the homecage, and direct contact was permitted. Social-interaction time was defined as sniffing, close social interaction and was scored using homemade MATLAB software by an observer blinded to group assignment.

### Novel-object recognition test

The novel-object recognition test was performed in the same arena used for the open-field test. During the familiarization phase, two identical cylindrical objects were positioned diagonally across the arena, and each mouse was allowed to freely explore the arena for 10 min. After a 5-7 h retention interval, one of the cylindrical objects was replaced with a novel triangular-prism-shaped object, while the position of the remaining familiar object was unchanged. Mice were then returned to the arena for the test phase. Object exploration was quantified as the time during which the center of the mouse body was located within 7 cm of the center of each object. Exploration times for the familiar and novel objects were subsequently compared within each mouse.

### EEG/EMG electrode implantation and EEG/EMG acquisition

After behavioral testing, mice underwent EEG/EMG electrode implantation at P90. Mice were anesthetized with isoflurane and placed in a stereotaxic frame. Following exposure of the skull, recording screws were positioned bilaterally over the dorsal hippocampus (anteroposterior-2.06 mm and mediolateral ±1.80 mm relative to bregma). the ground electrode and the reference screw were placed over the cerebellum. Two flexible EMG leads were inserted into the left and right nuchal muscles. The connector, screws, and wires were secured using dental cement. Animals recovered for 7 days for EEG/EMG recording. Following recovery, mice were connected to the recording apparatus and habituated to the tether and recording chamber for 2 days. Continuous EEG and EMG were then acquired for 24 hours beginning at zeitgeber time 0. Signals were recorded using Medusa Small Animal Electrophysiology Recording System, Bio-Signal technologies, at a sampling rate of 250 Hz, with line-noise filtering at 50 Hz.

### Sleep-stage classification and spectral analysis

Recordings were divided into 4-second epochs and classified as wakefulness, non-rapid-eye-movement (NREM) sleep, or rapid-eye-movement (REM) sleep using IntelliSleepScorer-v.1.2. Wakefulness was identified by low-amplitude, mixed-frequency EEG accompanied by variable or sustained EMG activity. NREM sleep was characterized by increased slow-wave activity and reduced EMG tone, whereas REM sleep was characterized by theta-dominant EEG and minimal nuchal EMG activity. Automated classifications were manually reviewed by an investigator blinded to treatment. The percentage of valid recording time spent in wakefulness, NREM sleep, and REM sleep was calculated separately for the 12-hour light and dark periods. Power spectral density was estimated using Welch’s method with a 256-point window, 128-point overlap, and 256-point fast Fourier transform. Frequency bands were defined as delta, 1–4 Hz; theta, 4–8 Hz; alpha, 8–12 Hz; sigma, 12–15 Hz; beta, 15–30 Hz; and gamma, 30–45 Hz. Band-limited RMS amplitudes were calculated for each vigilance state and photoperiod. Gamma-to-delta activity during NREM sleep, theta-to-delta activity during REM sleep, and alpha-to-delta activity during wakefulness were calculated from power amplitude. Mouse-level values, rather than individual epochs, were used for group comparisons.

### EEG analysis during social interaction

EEG and video recordings were synchronized during social interaction. Sniffing-associated social-contact bouts were defined as periods during which the subject sniffed the perianal region of the stimulus mouse. Annotation was performed using custom MATLAB software by an observer blinded to genotype and treatment. Bouts shorter than 3 s were excluded. For each bout, total EEG power and absolute power in the delta, theta, alpha, sigma, beta, and gamma bands were calculated using the spectral procedure described above. Relative power was expressed as band power divided by total power from 1 to 45 Hz. Theta-to-delta, alpha-to-delta, sigma-to-delta, beta-to-delta, and gamma-to-delta ratios were also calculated. Because animals contributed unequal numbers of social-contact bouts, individual bouts were not treated as independent biological replicates. Bout-level measurements were averaged within each mouse, and the mouse was used as the experimental unit.

### Tissue preparation and PV immunostaining

At P130, mice from the mMEF2C cohort were deeply anesthetized with isoflurane and perfused with ice-cold phosphate-buffered saline followed by 4% paraformaldehyde. Brains were removed, postfixed in 4% PFA overnight at 4°C. Brains were embedded in agarose and sectioned coronally at 50 μm thickness on a vibratome. Sections were sampled at bregma 2.58 - bregma-5.68 throughout the brain. Free-floating sections were blocked in 5% donkey serum albumin and 0.1% Triton X-100 for 90 minutes and incubated overnight at 4°C with rabbit monoclonal anti-PV antibody (Abcam ab181086, RRID AB_2313773, 1:1,000). Following washes, sections were incubated with secondary antibody (Thermo Fisher Scientific A-31572, RRID: AB_162543, 1:1000, Donkey-Anti-Rabbit Cy5), for 2 hours at room temperature. Nuclei were counterstained with DAPI. Sections from different groups were distributed across staining batches, and identical reagent and imaging settings were used within each batch.

### Whole-brain imaging and regional PV quantification

Coronal sections were scanned using an Olympus VS200 slide-scanning system with a 10 X objective. PV-immunoreactive profiles were segmented from the PV channel using Cellpose. The corresponding DAPI images were registered to the Allen Mouse Brain Common Coordinate Framework version 3 (CCFv3) using QuickNII. Segmented PV-immunoreactive profiles were assigned to atlas regions, and regional density was calculated as the number of segmented profiles divided by the analyzed regional area in square millimeters. Values from all brain regions were first combined within each mouse, after which mouse-level values were used for statistical analysis. Segmentation, registration quality control, and regional extraction were conducted with the analyst blinded to group assignment.

## Statistical analysis

Statistical analyses were performed using the Python statsmodels package and annotated using the statannotations package. The individual mouse was considered the biological replicate. Data are presented as mean ± SEM together with individual mouse-level observations. All tests were two-sided, and adjusted P < 0.05 was considered statistically significant. Effect sizes and 95% confidence intervals were reported where applicable. Single continuous outcomes involving three groups were analyzed using one-way ANOVA followed by Tukey’s post-hoc test. Three-chamber social-interaction and novel object data were analyzed using two-tailed paired t-tests to compare social versus empty stimuli during the social-approach phase and novel versus familiar stimuli during the novelty phase.

## Results

### Identification of brain-and muscle-enriched MEF2C isoforms

Transcript-level analysis across human tissues showed that ENST00000504921.7 was preferentially expressed across brain regions, whereas ENST00000340208.9 was most abundant in skeletal muscle (Supplementary Fig. S1A, B). We designated the encoded proteins nMEF2C and mMEF2C, respectively. nMEF2C corresponded to UniProt Q06413-1 (473 amino acids) and contained the alternatively spliced α1 region and the eight-residue β segment (SEDVDLLL). mMEF2C corresponded to Q06413-5 (483 amino acids), contained α2, and lacked β. Both retained the conserved MADS-box and MEF2 domains (Supplementary Fig. S1C, Fig. 1A).

**Figure 1.**
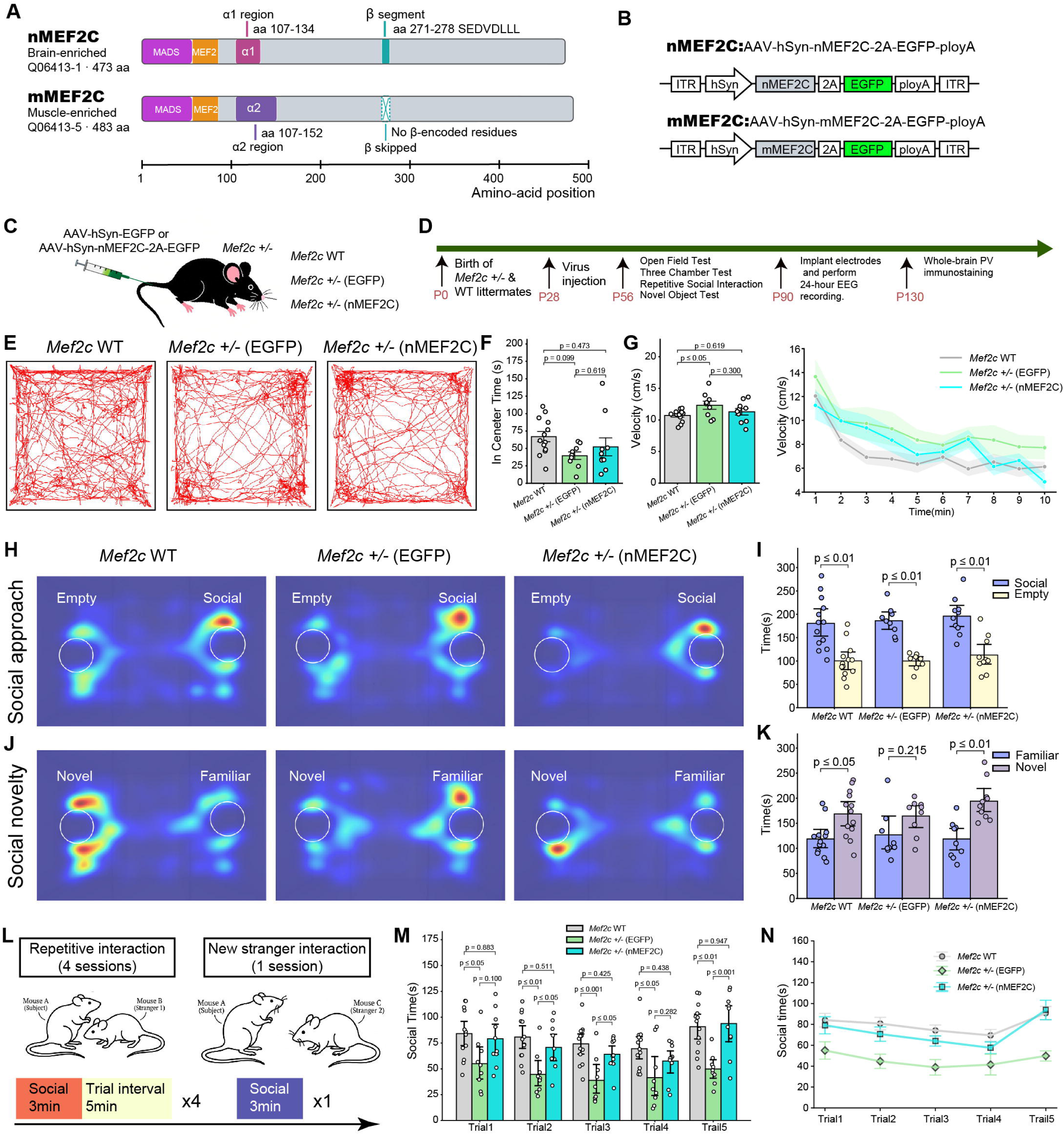
Human MEF2C isoforms, AAV gene-replacement vectors, and behavioral outcomes following juvenile nMEF2C delivery. (A) Protein schematics of the brain-enriched nMEF2C isoform (Q06413-1; 473 aa) and muscle-enriched mMEF2C isoform (Q06413-5; 483 aa). Both isoforms contain the conserved MADS-box and MEF2 DNA-binding domains. nMEF2C contains the α1 region (aa 107–134) and β-encoded segment (aa 271–278; SEDVDLLL), whereas mMEF2C contains the mutually exclusive α2 region (aa 107–152) and lacks β-encoded residues. (B) Schematics of the AAV transfer cassettes expressing nMEF2C or mMEF2C under the human synapsin promoter. Each MEF2C coding sequence was linked to EGFP through a T2A sequence and followed by a polyadenylation signal; the expression cassettes were flanked by AAV inverted terminal repeats. Schematics are not drawn to scale. (C) Experimental groups used to evaluate nMEF2C gene replacement: wild-type littermates, *Mef2c*^+/−^ mice receiving control AAV-hSyn-EGFP, and *Mef2c*^+/−^ mice receiving AAV-hSyn-nMEF2C-T2A-EGFP. (D) Experimental timeline. Viral vectors were administered at postnatal day 28 (P28). Open-field, three-chamber, repetitive social-interaction tests, and novel object tests were conducted beginning at P56. Electrode implantation and 24-h EEG/EMG recording were subsequently performed, followed by whole-brain PV immunostaining at P130. (E) Representative locomotor trajectories during the 10-min open-field test. (F) Time spent in the center of the open-field arena. (G) Mean locomotor velocity and minute-by-minute velocity across the 10-min test. Representative occupancy heatmaps during the social-approach phase (H) and social-novelty phase of the three-chamber test (J). Time spent interacting with the social stimulus versus the empty enclosure (I) and with the familiar versus novel social stimulus (K). All groups preferred the social stimulus to the empty enclosure. Wild-type and nMEF2C-treated *Mef2c*^+/−^ mice, but not EGFP-treated *Mef2c*^+/−^ mice, showed a significant preference for the novel social stimulus. (L) Repetitive social-interaction paradigm consisting of four 3-min encounters with the same stranger, separated by 5-min intervals, followed by a 3-min encounter with a new stranger. (M) Social-interaction time during each of the five trials. (N) Longitudinal representation of interaction time across the repeated and new-stranger trials. Bars and line plots show mean ± SEM; shaded areas in N indicate SEM, and circles represent individual mice (wild-type, n = 13; EGFP-treated *Mef2c*^+/−^, n = 9; nMEF2C- treated *Mef2c*^+/−^, n = 10). Comparisons are indicated by brackets, with exact *P* values or significance thresholds shown. (One-way ANOVA with Tukey’s post-hoc test was used for Fig. F, G, and M, and a two-tailed paired t-test was used for Fig. I, K.) aa, amino acids; AAV, adeno-associated virus; EGFP, enhanced green fluorescent protein; hSyn, human synapsin promoter; ITR, inverted terminal repeat; PV, parvalbumin; WT, wild type.

### Generation of *Mef2c*-haploinsufficient mice and MEF2C-expressing vectors

We generated mice carrying a heterozygous deletion of *Mef2c* exon 4 (*Mef2c*^+/ΔEx4^; hereafter *Mef2c*^+/−^; Supplementary Fig. S2A). For neuron-directed replacement, human nMEF2C or mMEF2C was expressed from the human synapsin promoter and linked to EGFP by T2A. Each cassette contained a Kozak sequence and human growth hormone polyadenylation signal and was flanked by AAV2 inverted terminal repeats (Fig. 1B).

To assess the spatial distribution of reporter expression, we performed GFP immunostaining approximately 100 days after vector administration. GFP-positive cells were broadly distributed across cortical, hippocampal, and subcortical regions (Supplementary Fig. S2B), demonstrating widespread reporter expression at the administered dose.

### Juvenile nMEF2C replacement selectively improves social phenotypes

*Mef2c*^+/−^ mice received AAV-hSyn-nMEF2C-T2A-EGFP or AAV-hSyn-EGFP at P28; wild-type littermates served as reference controls (Fig. 1C, D). At P56, control-treated mutants had higher open-field velocity than wild-type mice (*P* ≤ 0.05), whereas nMEF2C-treated mice differed from neither group (*P* = 0.619 versus wild type; *P* = 0.300 versus EGFP). Center time did not differ among groups (Fig. 1E–G).

All groups preferred the social stimulus to the empty enclosure, indicating preserved sociability. During the novelty phase, wild-type and nMEF2C-treated mice preferred the novel mouse (*P* ≤ 0.05 and *P* ≤ 0.01, respectively), whereas EGFP-treated mutants did not (*P* = 0.215; Fig. 1H-K). In repetitive social-interaction testing, nMEF2C treatment increased interaction relative to EGFP treatment during trials 2 and 3 and after introduction of a new partner (Fig. 1L–N). Thus, nMEF2C selectively improved novelty-related social measures. Complementary analyses of mean interaction time showed a similar pattern, supporting the social behavioral findings described above (Supplementary Fig. S3A–C). In the novel-object recognition test, all three groups preferentially investigated the novel object over the familiar object (Supplementary Fig. S3D, E). Thus, EGFP-treated *Mef2c*^+/−^ mice exhibited impaired social-novelty responses despite preserved novel-object preference. This pattern is consistent with a possible deficit in social recognition memory.

### nMEF2C replacement modifies sleep–wake architecture and selected EEG features

After recovery and habituation following electrode implantation, mice underwent continuous EEG/EMG recording for 24 h (Fig. 2A, B). During the dark phase, EGFP-treated *Mef2c*^+/−^ mice showed less wakefulness and more NREM and REM sleep than wild-type mice (*P* ≤ 0.05 for each; Fig. 2C). nMEF2C treatment increased wakefulness and reduced NREM sleep relative to EGFP treatment (*P* ≤ 0.05), yielding values comparable to wild type. REM sleep shifted toward wild type but did not differ from EGFP treatment (*P* = 0.116). During the light phase, wakefulness and NREM sleep were unchanged; the reduced REM proportion in EGFP-treated mutants was intermediate after nMEF2C treatment, without a significant direct treatment effect (Fig. 2D).

**Figure 2.**
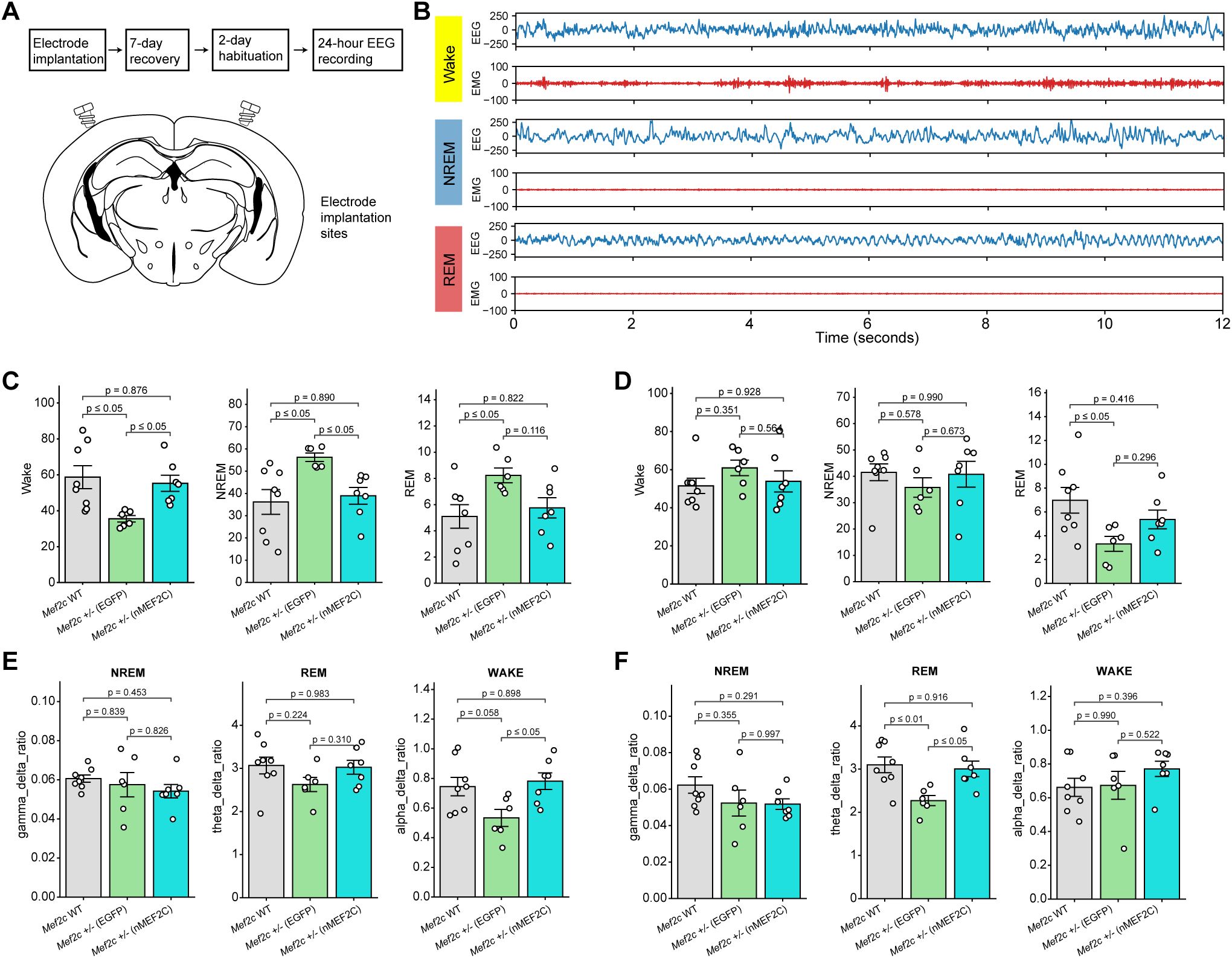
AAV-nMEF2C gene replacement modifies sleep–wake architecture and selected state-dependent EEG features in *Mef2c^+/−^* mice. (A) Experimental workflow for continuous EEG/EMG recording. Following electrode implantation, mice underwent a 7-day recovery period and 2 days of habituation before 24-h recording. (B) Representative 12-s EEG and EMG traces illustrating wakefulness, non-rapid eye movement (NREM) sleep, and rapid eye movement (REM) sleep. (C) Proportions of time spent awake or in NREM and REM sleep during the dark phase. Compared with wild-type mice, EGFP-treated *Mef2c*^+/−^ mice exhibited reduced wakefulness and increased NREM and REM sleep. nMEF2C treatment increased wakefulness and decreased NREM sleep relative to EGFP treatment, resulting in values comparable to those of wild-type mice. The REM proportion shifted toward the wild-type level but did not differ significantly between the two mutant treatment groups. (D) Proportions of wakefulness, NREM sleep, and REM sleep during the light phase. Wakefulness and NREM sleep did not differ significantly among groups. EGFP-treated *Mef2c*^+/−^ mice exhibited a lower REM proportion than wild-type mice, whereas the nMEF2C-treated group showed an intermediate value that did not differ significantly from either comparison group. (E) State-dependent EEG band-power ratios during the dark phase, including the γ/δ ratio during NREM sleep, θ/δ ratio during REM sleep, and α/δ ratio during wakefulness. The NREM γ/δ and REM θ/δ ratios did not differ significantly among groups. The wake α/δ ratio was higher in nMEF2C-treated mice than in EGFP-treated mutants and was comparable to that in wild-type mice; the wild-type versus EGFP-treated comparison showed a nonsignificant trend. (F) Corresponding EEG ratios during the light phase. The NREM γ/δ and wake α/δ ratios did not differ significantly among groups. The REM θ/δ ratio was reduced in EGFP-treated *Mef2c*^+/−^ mice relative to wild-type mice and increased following nMEF2C treatment to a level comparable to that of wild-type controls. Bars show mean ± SEM, and each circle represents one mouse (wild-type, n = 8; EGFP-treated *Mef2c*^+/−^, n = 6; nMEF2C-treated *Mef2c*^+/−^, n = 7). Data were analyzed by one-way ANOVA followed by Tukey’s multiple-comparisons test. Comparisons are indicated by brackets, with exact *P* values or significance thresholds shown. EEG, electroencephalography; EMG, electromyography; NREM, non-rapid eye movement; REM, rapid eye movement.

Spectral effects were state and phase specific. Dark-phase wake alpha-to-delta ratio showed a trend toward reduction in EGFP-treated mutants versus wild type (*P* = 0.058) and increased after nMEF2C treatment (*P* ≤ 0.05; Fig. 2E). During the light phase, the REM theta-to-delta ratio was reduced in EGFP-treated mutants (*P* ≤ 0.01) and increased after nMEF2C treatment (*P* ≤ 0.05; Fig. 2F). Other tested ratios were unchanged. Absolute band RMS amplitudes were also unchanged during the dark phase. Light-phase NREM delta and theta amplitudes were elevated in EGFP-treated mutants, but nMEF2C did not differ significantly from EGFP treatment (Supplementary Fig. S4).

### nMEF2C replacement tends to normalize elevated sigma activity during social interaction

To examine neural activity associated with social behavior, we analyzed EEG spectral features during annotated sniffing episodes. When sigma activity was separated from the broader beta-frequency range, EGFP-treated *Mef2c*^+/−^ mice exhibited increased absolute sigma power, relative sigma power, and the sigma-to-delta ratio compared with wild-type mice (P ≤ 0.01, P ≤ 0.05, and P ≤ 0.05, respectively; Supplementary Fig. S5B-C). In nMEF2C-treated *Mef2c*^+/−^ mice, all three sigma measures shifted toward wild-type values and no longer differed significantly from wild-type controls. Absolute and relative sigma power tended to be lower than in EGFP-treated mutants, although the direct treatment comparisons did not reach statistical significance (P = 0.061 and P = 0.063, respectively). The sigma-to-delta ratio also showed an intermediate value after treatment, but did not differ significantly between the two mutant groups (P = 0.309).

Most other absolute and relative band-power measures and spectral ratios were unchanged. Absolute gamma power was higher in nMEF2C-treated mice than in wild-type mice (P ≤ 0.05), but did not differ from that in EGFP-treated mutants (P = 0.778), and therefore did not represent correction of a mutant-associated phenotype. Collectively, these exploratory findings identify increased sigma activity during social interaction as an electrophysiological feature of *Mef2c* haploinsufficiency and suggest directional, but not statistically established, normalization following nMEF2C gene replacement.

### mMEF2C replacement selectively improves social-novelty phenotypes

In a separate cohort, EGFP-treated *Mef2c*^+/−^ mice had higher open-field velocity than wild-type mice (*P* ≤ 0.001; Fig. 3A–C). mMEF2C-treated mice were intermediate but differed from neither group (*P* = 0.054 versus wild type; *P* = 0.057 versus EGFP). Both mutant groups spent less time in the center than wild-type mice, with no treatment effect (Fig. 3D).

**Figure 3.**
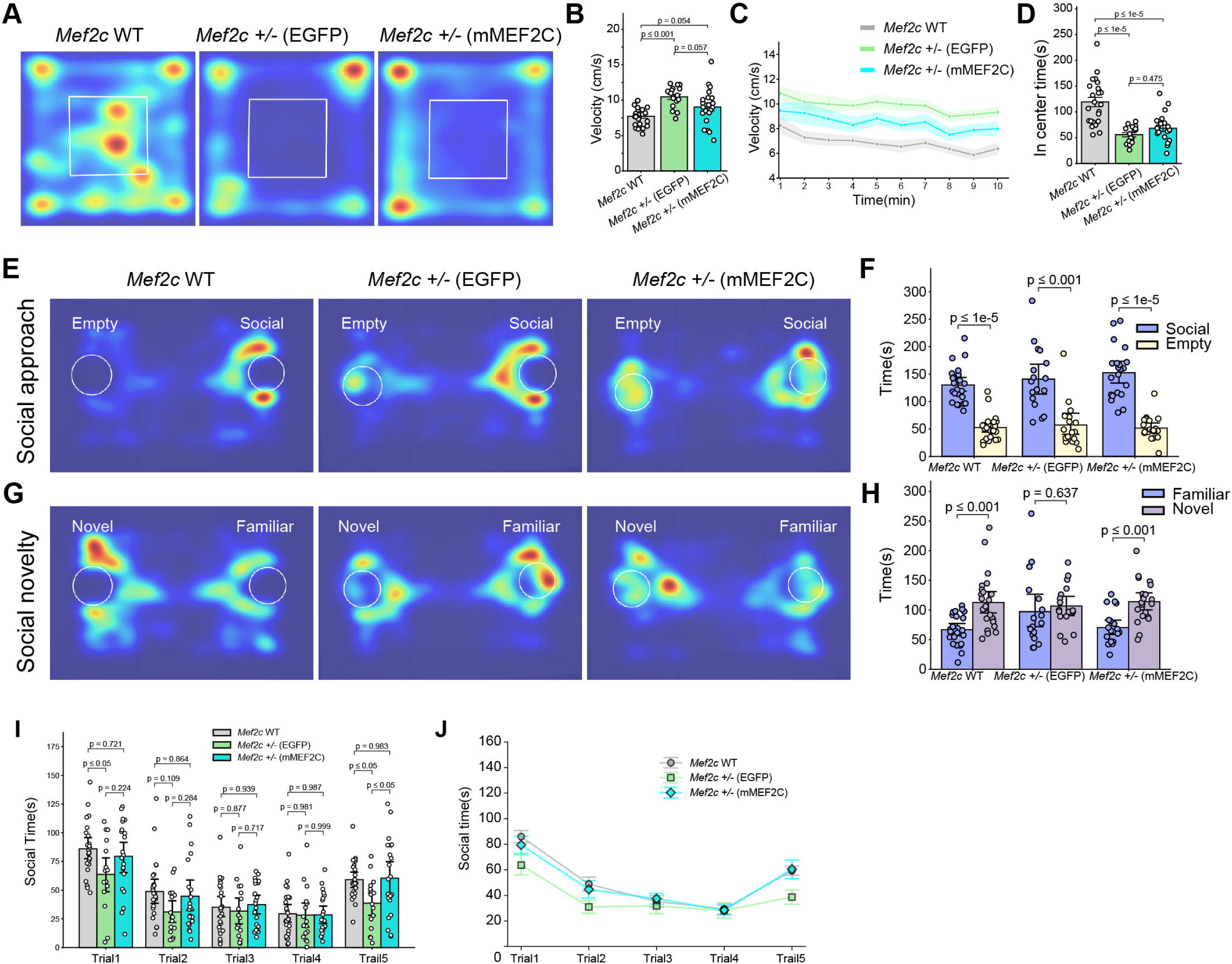
Behavioral outcomes following AAV-mMEF2C gene replacement in *Mef2c^+/−^* mice. (A) Representative occupancy heatmaps from the 10-min open-field test for wild-type mice, EGFP-treated *Mef2c*^+/−^ mice, and mMEF2C-treated *Mef2c*^+/−^ mice. The white square indicates the predefined center region. (B) Mean locomotor velocity during the open-field test. EGFP-treated *Mef2c*^+/−^ mice exhibited greater velocity than wild-type mice. The mean velocity of mMEF2C-treated mice was intermediate and did not differ significantly from either wild-type or EGFP-treated mutant mice. (C) Minute-by-minute locomotor velocity across the 10-min test. (D) Time spent in the center region. Both EGFP-and mMEF2C-treated *Mef2c*^+/−^ mice spent less time in the center than wild-type mice, with no significant difference between the two mutant treatment groups. (E) Representative occupancy heatmaps during the social-approach phase of the three-chamber test. (F) Time spent interacting with the social stimulus or empty enclosure. All three groups showed a significant within-group preference for the social stimulus. (G) Representative occupancy heatmaps during the social-novelty phase. (H) Time spent interacting with the familiar or novel social stimulus. Wild-type and mMEF2C-treated *Mef2c*^+/−^ mice showed a significant within-group preference for the novel stimulus, whereas EGFP-treated *Mef2c*^+/−^ mice did not. (I) Social-interaction time during four repeated encounters with the same stranger (trials 1–4) and a subsequent encounter with a new stranger (trial 5). EGFP-treated *Mef2c*^+/−^ mice interacted less than wild-type mice during trials 1 and 5. During trial 5, mMEF2C-treated mice interacted significantly more than EGFP-treated mutants and did not differ from wild-type mice. No significant treatment effects were detected during trials 2–4. (J) Longitudinal representation of social-interaction time across the five trials. Bars and line plots show mean ± SEM; shaded areas in J indicate SEM, and circles represent individual mice (wild-type, n = 24; EGFP-treated *Mef2c*^+/−^, n = 17; mMEF2C-treated *Mef2c*^+/−^, n = 25). Between-group comparisons in B, D, and I were performed using one-way ANOVA followed by Tukey’s multiple-comparisons test. Within-group stimulus comparisons in F and H were performed using two-tailed paired *t*-tests. Comparisons are indicated by brackets, with exact *P* values or significance thresholds shown. AAV, adeno-associated virus; EGFP, enhanced green fluorescent protein; WT, wild type.

Baseline sociability was preserved in all groups (Fig. 3E, F). Wild-type and mMEF2C-treated mice preferred the novel social partner (*P* ≤ 0.001 for each), whereas EGFP-treated mutants did not (*P* = 0.637; Fig. 3G, H). In repetitive interaction testing, mMEF2C treatment did not alter trials 1–4 but increased interaction with the new partner in trial 5 relative to EGFP treatment (*P* ≤ 0.05; Fig. 3I, J). These findings support a selective improvement in social-novelty responsiveness. Because the nMEF2C and mMEF2C experiments used separate cohorts, they do not establish relative isoform efficacy. Complementary analyses of mean interaction time were consistent with the primary social-behavior analyses, supporting preserved baseline sociability and the observed patterns of social-novelty preference across groups (Supplementary Fig. S6A– C). Consistent with findings in the nMEF2C cohort, wild-type, EGFP-treated *Mef2c*^+/−^, and mMEF2C-treated *Mef2c*^+/−^ mice all preferentially investigated the novel object over the familiar object in the novel-object recognition test (Supplementary Fig. S6D, E).

### MEF2C replacement produces region-selective shifts in PV-immunoreactive phenotypes

PV-immunoreactive profiles were segmented from serial coronal sections and registered to the Allen Mouse Brain Common Coordinate Framework for atlas-based quantification (Fig. 4A). In the nMEF2C cohort, EGFP-treated mutants had lower PV-immunoreactive profile densities than wild-type mice in MOp, MOs, SSp, SSs, RSP, VIS, AUD, and CA3 (Fig. 4B, upper). nMEF2C significantly increased profile density relative to EGFP treatment in MOs, the only region meeting the conservative criterion of a mutant abnormality plus a direct corrective treatment effect. Other cortical means shifted toward wild type without significant direct treatment comparisons, and SSp and CA3 remained below wild type.

**Figure 4.**
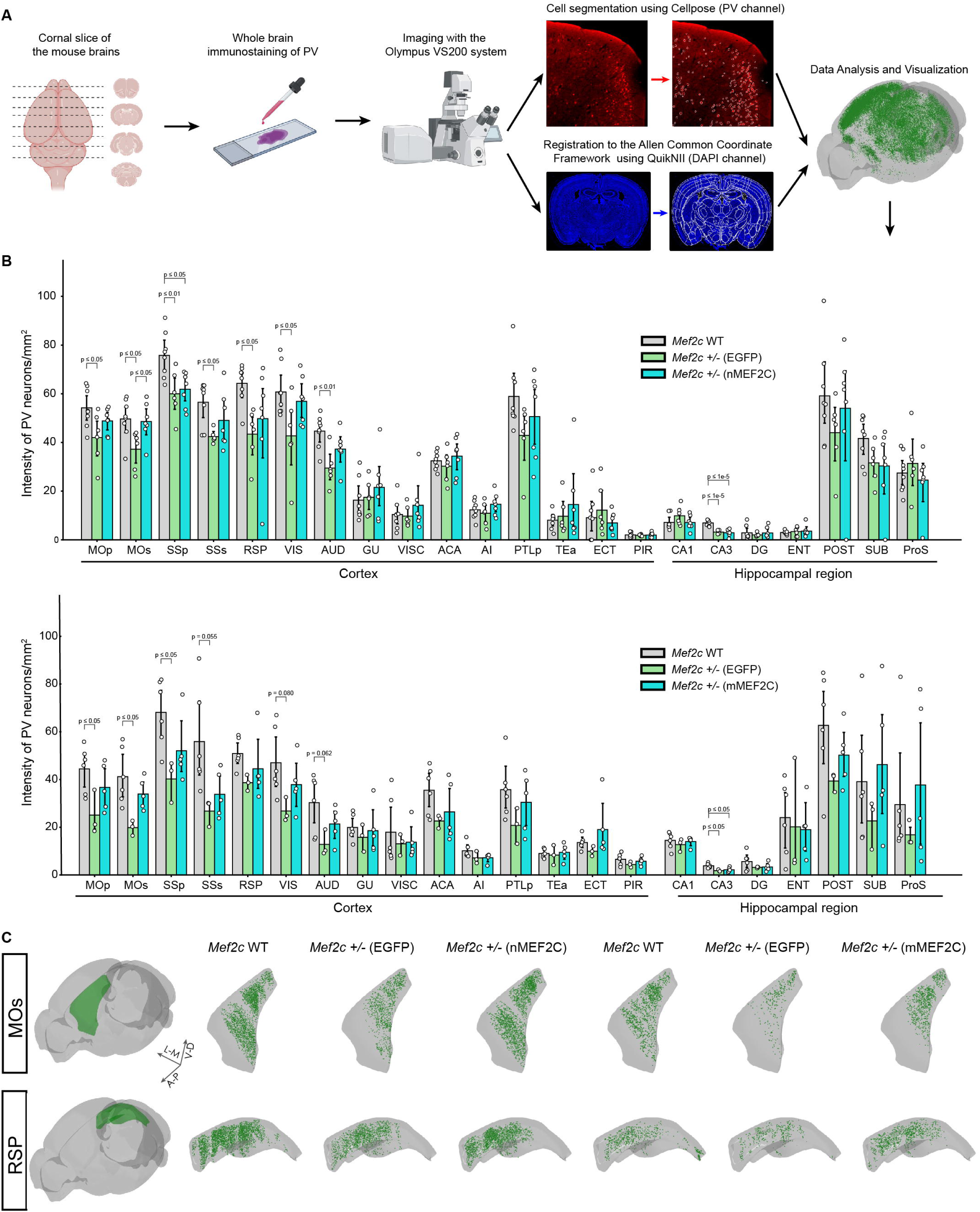
Region-selective changes in PV-immunoreactive profiles following nMEF2C or mMEF2C gene replacement in *Mef2c^+/^*^−^ mice. (A) Workflow for atlas-based whole-brain analysis of parvalbumin (PV) immunoreactivity. Serial coronal sections were immunostained for PV and imaged using an Olympus VS200 slide scanner. PV-immunoreactive profiles were segmented from the PV channel using Cellpose, while corresponding DAPI images were registered to the Allen Mouse Brain Common Coordinate Framework using QuickNII. Segmented profiles were subsequently assigned to anatomical regions for quantification and visualization. (B) Regional densities of PV-immunoreactive profiles in independent nMEF2C (upper) and mMEF2C (lower) treatment cohorts. Groups comprised wild-type mice, EGFP-treated *Mef2c^+/−^* mice and *Mef2c^+/−^* mice treated with the indicated MEF2C isoform. In the nMEF2C cohort, EGFP-treated mutants exhibited lower PV-positive profile densities than wild-type mice in several cortical regions and CA3. nMEF2C treatment increased profile density relative to EGFP treatment in the secondary motor area (MOs), whereas the reductions in the primary somatosensory area (SSp) and CA3 persisted relative to wild type. Other cortical regions showed directional shifts toward wild-type values without significant direct treatment effects. In the mMEF2C cohort, PV-positive profile densities similarly shifted toward wild-type values in several affected regions; however, no significant direct comparison between the EGFP-and mMEF2C-treated groups was detected. The two isoform cohorts were analyzed independently and were not statistically compared with each other, as the sampling fractions differed between them (one-quarter of sections for nMEF2C versus one-sixth for mMEF2C). (C) Representative atlas-registered distributions of segmented PV-immunoreactive profiles in MOs and the retrosplenial area (RSP). Green points indicate segmented PV-positive profiles within the regional boundaries shown in gray. Coordinates are shown as A-P (anteroposterior), D-V (dorsoventral), and M-L (mediolateral). Each circle in (B) represents one mouse, and bars indicate mean ± SEM. Pairwise *P* values are displayed above the corresponding brackets. Brain areas and abbreviations: Primary motor area: MOp; Secondary motor area: MOs; Primary somatosensory area: SSp; Supplemental somatosensory area: SSs; Retrosplenial area: RSP; Visual areas: VIS; Auditory areas: AUD; Gustatory areas: GU; Visceral area: VISC; Anterior cingulate area: ACA; Agranular insular area: AI; Posterior parietal association areas: PTLp; Temporal association areas: TEa; Ectorhinal area: ECT; Piriform area: PIR; Field CA1: CA1; Field CA3: CA3; Dentate gyrus: DG; Entorhinal area: ENT; POST: postsubiculum; Subiculum: SUB; Prosubiculum: ProS.

In the mMEF2C cohort, EGFP-treated mutants showed reductions in MOp, MOs, SSp, SSs (*P* =0.055), VIS (*P* =0.08), AUD (*P*=0.062), and CA3 (Fig. 4B, lower). Several means shifted toward wild type after mMEF2C treatment, but no region showed a significant direct EGFP-versus-mMEF2C effect; CA3 remained below wild type. Thus, PV-associated phenotypes and their response to MEF2C replacement were anatomically selective.

## Discussion

Neuron-directed MEF2C replacement initiated at P28 produced selective improvements in *Mef2c*-haploinsufficient mice. Both isoforms improved novelty-related social outcomes, whereas open-field effects were limited. nMEF2C additionally corrected dark-phase wakefulness and NREM sleep abnormalities and modified selected EEG ratios. PV-immunoreactive abnormalities were region selective, with a statistically supported corrective treatment effect confined to the secondary motor area (MOs) after nMEF2C replacement. These findings support continued responsiveness of selected phenotypes to juvenile MEF2C supplementation.

These findings complement postnatal pharmacological rescue and mutation-specific DNA and RNA editing studies.^9,11,12^ Given the allelic heterogeneity of MCHS,^1–4^ delivery of a functional coding sequence could potentially address deletions and diverse loss-of-function variants. Here, gene replacement denotes exogenous MEF2C cDNA delivery to compensate for haploinsufficiency, not repair of the endogenous allele. Because treatment occurred during the juvenile period and no pretreatment measurements were obtained, the study does not establish reversal of pre-existing developmental abnormalities or define an adult therapeutic window.

The overlapping social effects of nMEF2C and mMEF2C suggest that both isoforms can support neuronal function despite their α/β splicing differences. However, the isoforms were tested in separate cohorts, equivalent transgene exposure was not demonstrated, and sleep and EEG outcomes were assessed only for nMEF2C. No efficacy ranking can therefore be made. The dissociation between novelty-related improvement and limited open-field effects also argues against a nonspecific increase in exploration.

The sleep results extend the known relationship between MEF2C and sleep-dependent cortical regulation.^10^ Their state-and phase-specific nature, together with largely unchanged absolute band amplitudes, argues against broad normalization of cortical oscillations. The dark-phase wake alpha-to-delta increase represents treatment-associated modulation because the baseline genotype contrast was not significant. During social sniffing, elevated sigma-range activity shifted toward wild-type values after nMEF2C treatment, but direct treatment comparisons were not significant. These exploratory changes therefore do not establish electrophysiological rescue or social specificity. Sleep/EEG and social improvements occurred at the group level; within-animal correlations and causal relationships remain untested.

The regional PV findings are compatible with the hypothesis that functional correction within selected circuits may support behavioral improvement without widespread anatomical normalization. Selimbeyoglu et al. showed that increasing medial prefrontal PV-interneuron excitability rescued social-interaction deficits in *Cntnap2*-deficient mice.^13^ Bicks et al. similarly found that activating dorsomedial prefrontal PV interneurons ameliorated social deficits following juvenile social isolation.^14^ These studies establish precedents for behavioral benefit through localized inhibitory-circuit modulation, but they do not demonstrate that MOs mediates the social effects of MEF2C replacement.

In our study, a significant corrective effect on PV-immunoreactive profile density was confined to MOs after nMEF2C treatment. Improved inhibitory regulation in this region could contribute to social investigation, but this remains a hypothesis. Social-novelty responses also improved after mMEF2C replacement without significant regional PV-density increases; the detected MOs change therefore cannot provide a complete explanation of the behavioral benefits. Moreover, profile density cannot distinguish altered PV expression or detectability from changes in interneuron abundance, and it does not establish restored inhibitory function. The MOs finding is consequently a candidate cellular correlate of treatment. Region-and cell-type-specific manipulations, vector–PV colocalization, and circuit recordings are needed to test causality.

Translation will require quantitative transgene-expression and target-engagement measurements, dose–response studies, a wild-type-plus-MEF2C safety group, and evaluation of biodistribution, durability, immunogenicity, neuropathology, sex-dependent responses, and long-term safety. Synapsin-driven expression does not reproduce endogenous temporal, activity-dependent, or cell-type-specific MEF2C regulation and may not address non-neuronal contributions.^7,8^ A clinical construct would also require removal of EGFP and evaluation of a suitable capsid, route, and manufacturing process. Subject to these limitations, the present results support continued development of MEF2C replacement as a potentially genotype-flexible therapeutic strategy.

## Acknowledgments

The authors are deeply grateful to MEF2C-related disorder families. Their participation, support, and commitment to advancing MEF2C research were invaluable to this study.

ChatGPT (OpenAI) was used to assist with language editing, manuscript organization, and drafting. The authors independently reviewed and verified all scientific content, analyses, interpretations, and references and accept full responsibility for the manuscript.

## Author Contributions

Z.J. and Z.Q. designed the study. Z.J., C.Y., and T.L. performed the behavioral tests, EEG recording and immunohistochemistry. Y. Yuan and J.W. carried out the virus injections. Z.J., C.Y., T.L., and Y.L. analyzed the behavioral, EEG, and immunohistochemistry data. C.Y., G.T., and Y. Yang assisted with atlas registration for PV neuron counting. Y.Y. and Y.Z. managed mouse breeding and genotyping. Z.Q. wrote the manuscript.

## Statements and declarations

### Data availability

The data supporting the findings of this study are available from the corresponding authors upon reasonable request.

