## supplemental figure 1-6 for "*Juvenile* AAV-Mediated MEF2C Gene Replacement Ameliorates Selected Phenotypes in *Mef2c*-Haploinsufficient Mice"

### Supplementary Figure Legends

**Supplementary Figure 1**

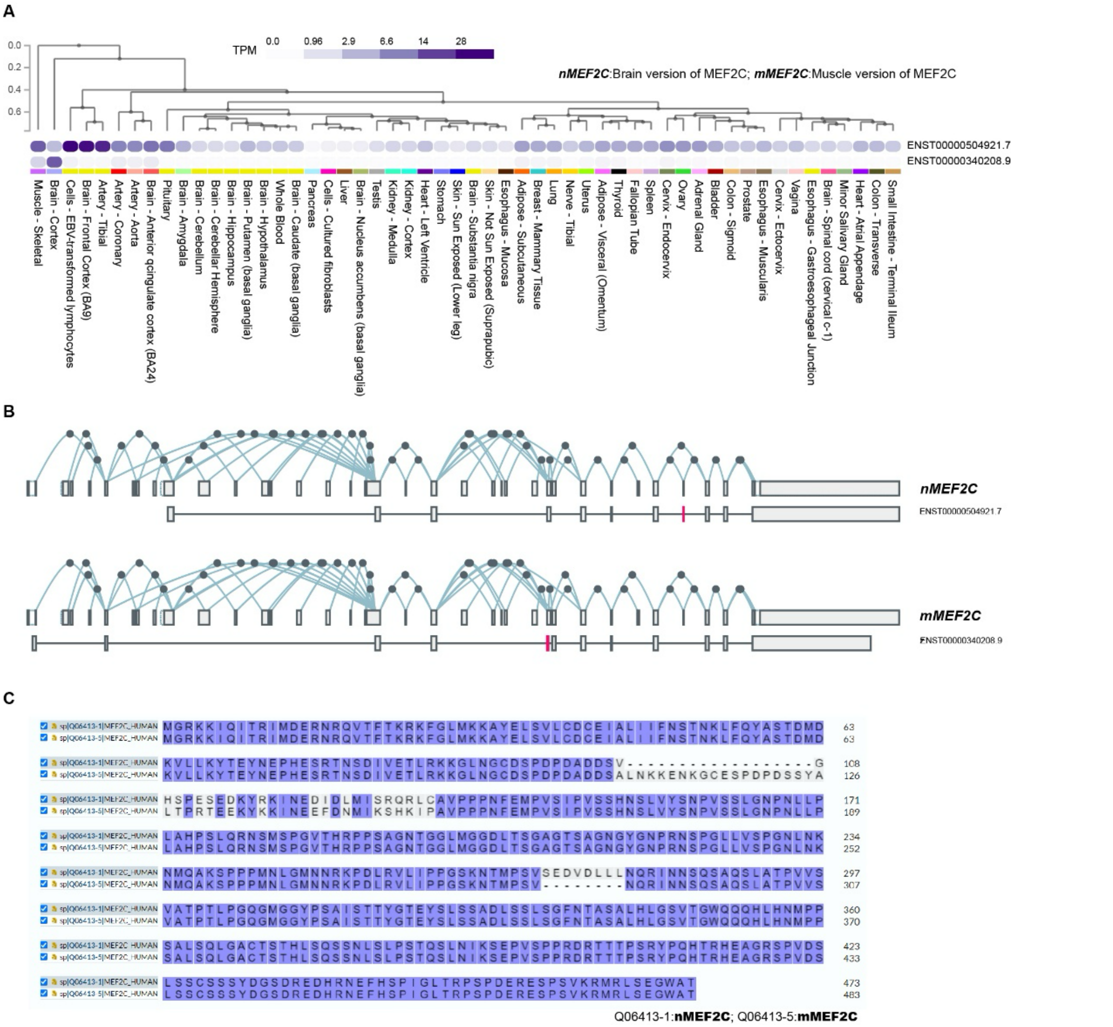


**Supplementary Figure 1. Tissue-enriched expression and alternative splicing of human MEF2C isoforms.**

(A) Transcript-level expression of two major human *MEF2C* transcripts across tissues in the GTEx human transcriptome dataset. Expression is presented as transcripts per million (TPM), with darker colors indicating higher expression. Tissues were hierarchically clustered according to their transcript-expression profiles. ENST00000504921.7 was preferentially expressed in brain tissues, whereas ENST00000340208.9 showed prominent expression in skeletal muscle. Based on these expression patterns, the corresponding protein isoforms were designated nMEF2C and mMEF2C, respectively, in this study. (B) Ensemble exon structures and splice-junction maps for ENST00000504921.7 and ENST00000340208.9. Exons are shown as boxes, introns as connecting lines, and annotated splice junctions as arcs. Isoform-distinguishing alternatively spliced segments are highlighted in red. (C) Pairwise amino-acid sequence alignment of the proteins encoded by the two transcripts: UniProt Q06413-1, corresponding to nMEF2C, and Q06413-5, corresponding to mMEF2C. The proteins share the conserved N-terminal DNA-binding regions but differ within the mutually exclusive α1/α2 region. Q06413-1 contains the α1 sequence and β-encoded segment, whereas Q06413-5 contains the longer α2 sequence and lacks the β-encoded residues, resulting in protein lengths of 473 and 483 amino acids, respectively. aa, amino acids; GTEx, Genotype-Tissue Expression; TPM, transcripts per million.

**Supplementary Figure 2**

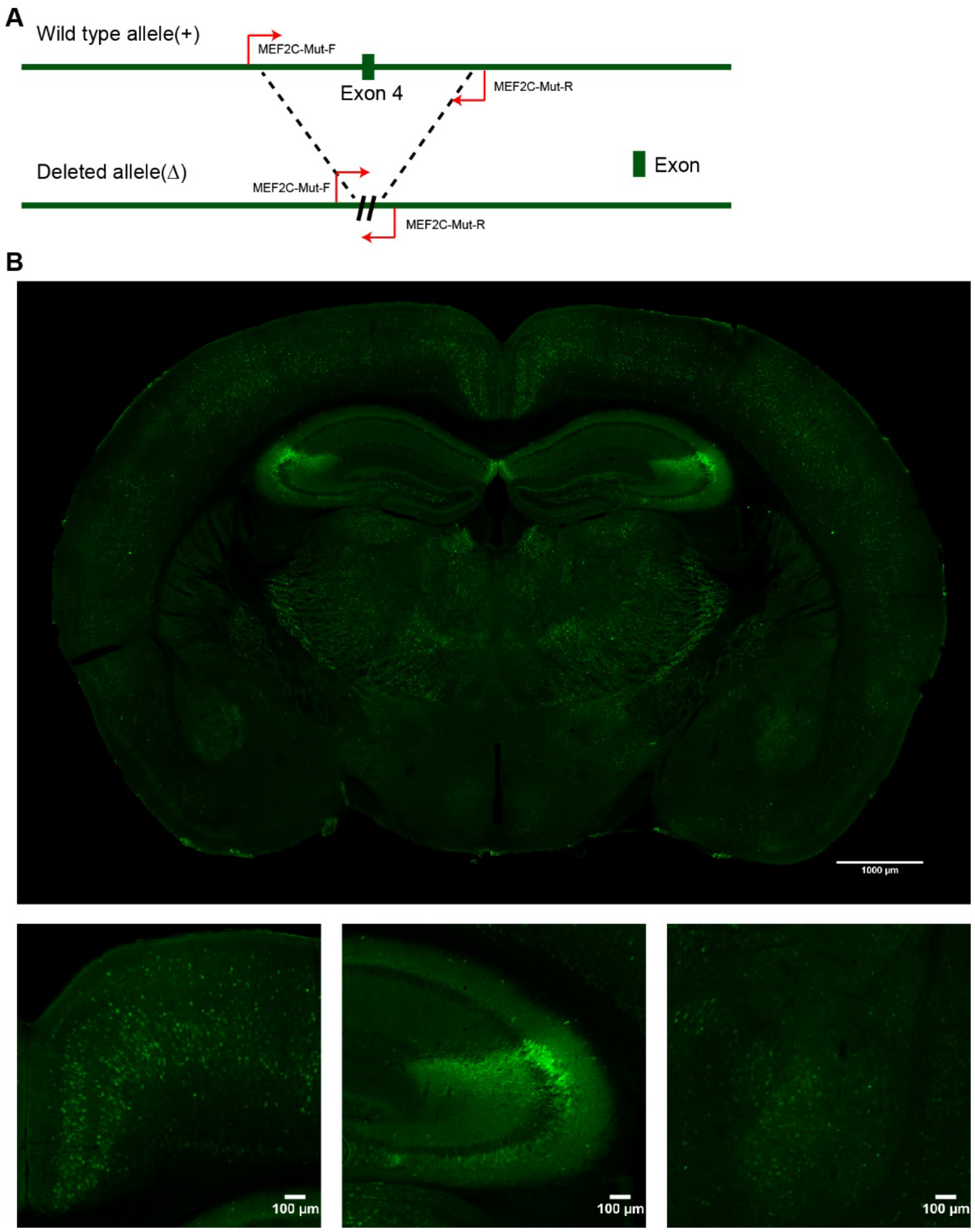


**Supplementary Figure 2. Generation and genotyping of *Mef2c* exon 4-deletion mice.**

(A) Schematic of the wild-type and exon 4-deleted *Mef2c* alleles. The wild-type allele contains exon 4, whereas the deleted allele lacks the exon 4-containing genomic segment indicated by the dashed lines. The positions and orientations of the genotyping primers, MEF2C-Mut-F and MEF2C-Mut-R, are shown in red. The schematic is not drawn to scale. (B) Representative EGFP immunostaining (green) approximately 100 days after AAV administration. The upper image shows a coronal brain section with GFP-positive cells distributed across cortical, hippocampal, and subcortical regions. Representative higher-magnification images of these regions are shown below (left to right). Scale bars: 1,000 µm (upper image) and 100 µm (lower images).

**Supplementary Figure 3**

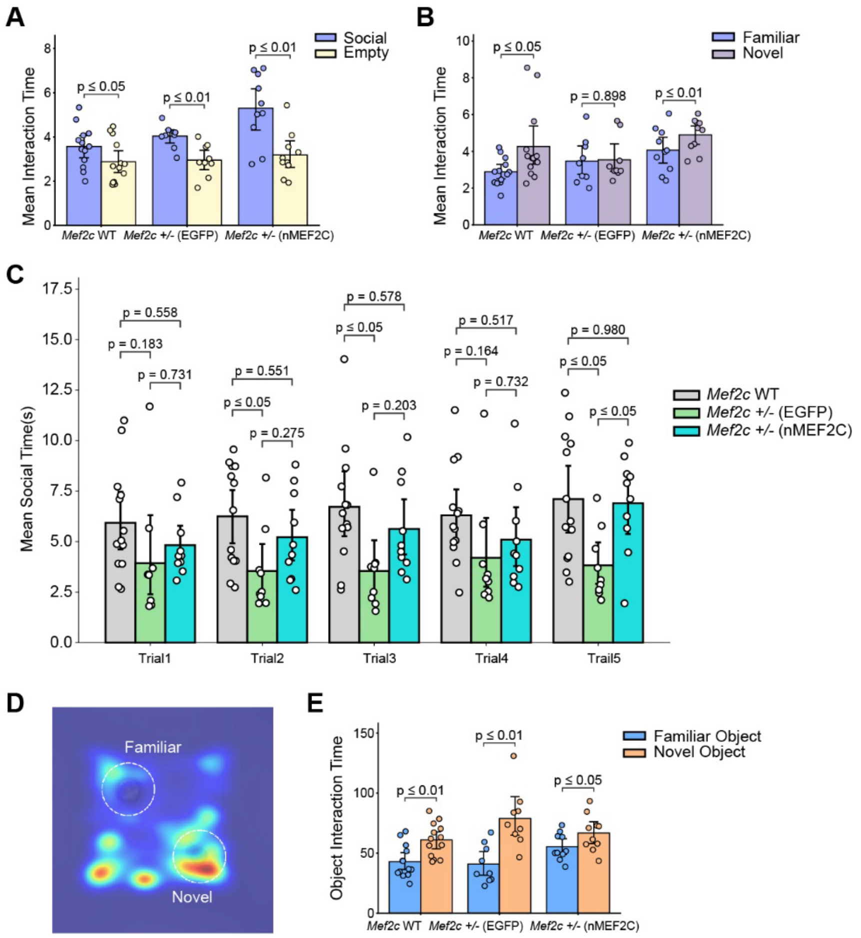


**Supplementary Figure 3. Additional social-interaction analyses and novel-object recognition in the nMEF2C cohort.**

(A) Mean interaction time with the social stimulus versus the empty enclosure during the social-approach phase of the three-chamber test. All three groups showed a significant preference for the social stimulus. (B) Mean interaction time with the familiar versus novel conspecific during the social-novelty phase. Wild-type and nMEF2C-treated *Mef2c*$^{+/-}$ mice showed significant novel-conspecific preference, whereas EGFP-treated *Mef2c*$^{+/-}$ mice did not. (C) Mean social-interaction time during four repeated encounters with the same intruder (trials 1–4), followed by an encounter with a novel intruder (trial 5). During trial 5, nMEF2C-treated *Mef2c*$^{+/-}$ mice showed significantly longer mean social-interaction time than EGFP-treated mutants. (D) Representative occupancy heatmap during the novel-object recognition test. Dashed circles indicate the familiar and novel object locations. (E) Time spent investigating the familiar and novel objects. All three groups showed a significant preference for the novel object. Groups comprised wild-type mice, EGFP-treated *Mef2c*$^{+/-}$ mice, and nMEF2C-treated *Mef2c*$^{+/-}$ mice. Bars indicate mean ± SEM, and individual points represent individual mice. Brackets indicate statistical comparisons, with exact P values or significance thresholds shown. EGFP, enhanced green fluorescent protein; WT, wild type.

**Supplementary Figure 4**

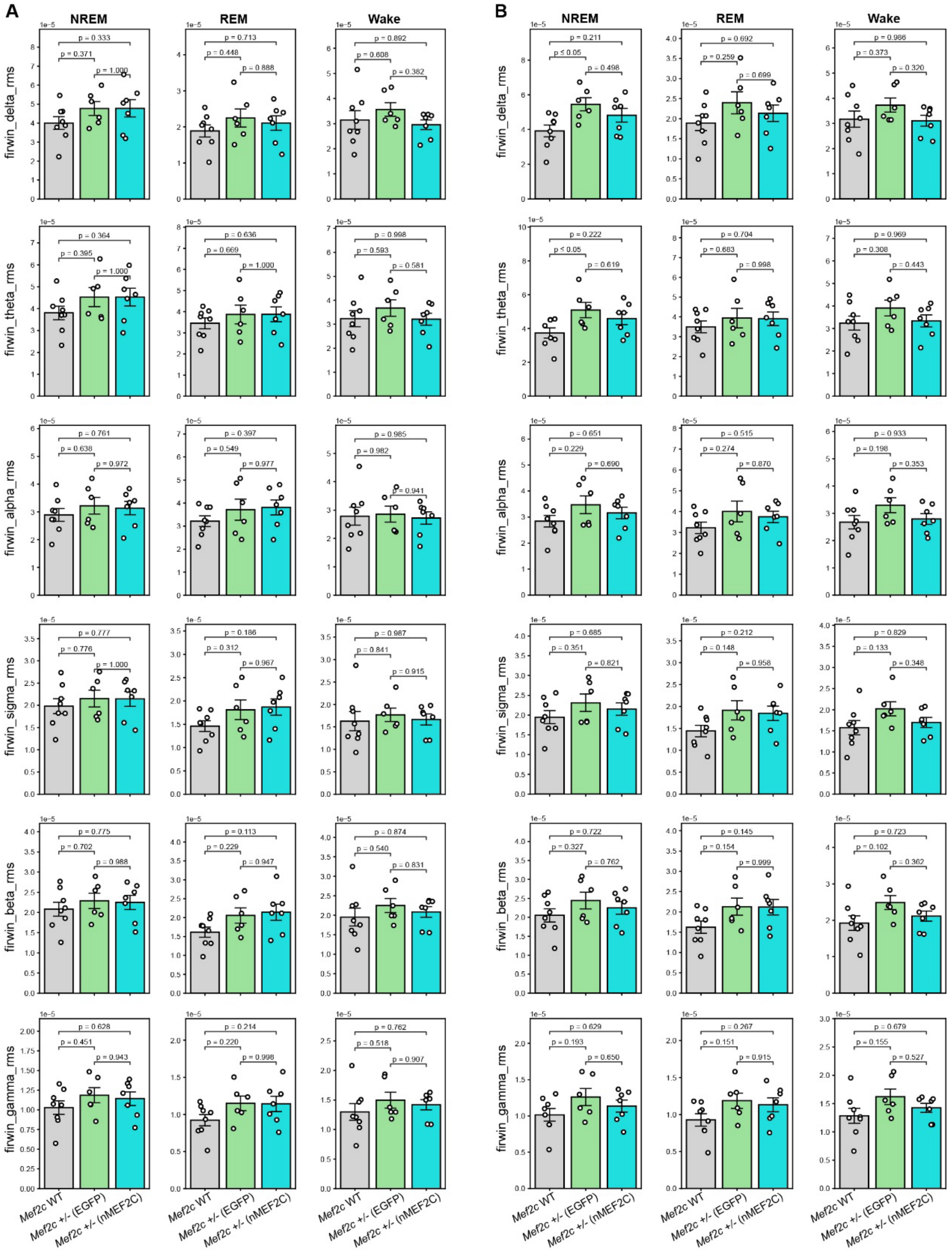


**Supplementary Figure 4. State-dependent absolute EEG band amplitudes following nMEF2C gene replacement.**

(A, B) Root-mean-square (RMS) amplitudes of finite-impulse-response-filtered EEG frequency bands were quantified separately during non-rapid eye movement (NREM) sleep, rapid eye movement (REM) sleep, and wakefulness. The six rows show, from top to bottom, δ-, θ-, α-, σ-, β- and γ-band RMS amplitudes. Frequency ranges used to define each band are provided in the Materials and Methods. (A) State-dependent EEG band amplitudes during the dark phase. No significant differences were detected among wild-type mice, EGFP-treated *Mef2c*$^{+/-}$ mice, and nMEF2C-treated *Mef2c*$^{+/-}$ mice for any frequency band or vigilance state. (B) Corresponding EEG band amplitudes during the light phase. EGFP-treated *Mef2c*$^{+/-}$ mice exhibited higher NREM δ- and θ-band RMS amplitudes than wild-type mice. Values in nMEF2C-treated mice were intermediate, but neither measure differed significantly between the EGFP- and nMEF2C-treated groups. No significant group differences were detected for the remaining frequency bands or vigilance states. Bars show mean ± SEM, and each circle represents one mouse (wild-type, n = 8; EGFP-treated *Mef2c*$^{+/-}$, n = 6; nMEF2C-treated *Mef2c*$^{+/-}$, n = 7). Data were analyzed separately for each band and vigilance state using one-way ANOVA followed by Tukey's multiple-comparisons test. Comparisons are indicated by brackets, with exact *P* values or significance thresholds shown. EEG, electroencephalography; NREM, non-rapid eye movement; REM, rapid eye movement; RMS, root mean square; WT, wild type.

### Supplementary figure 5

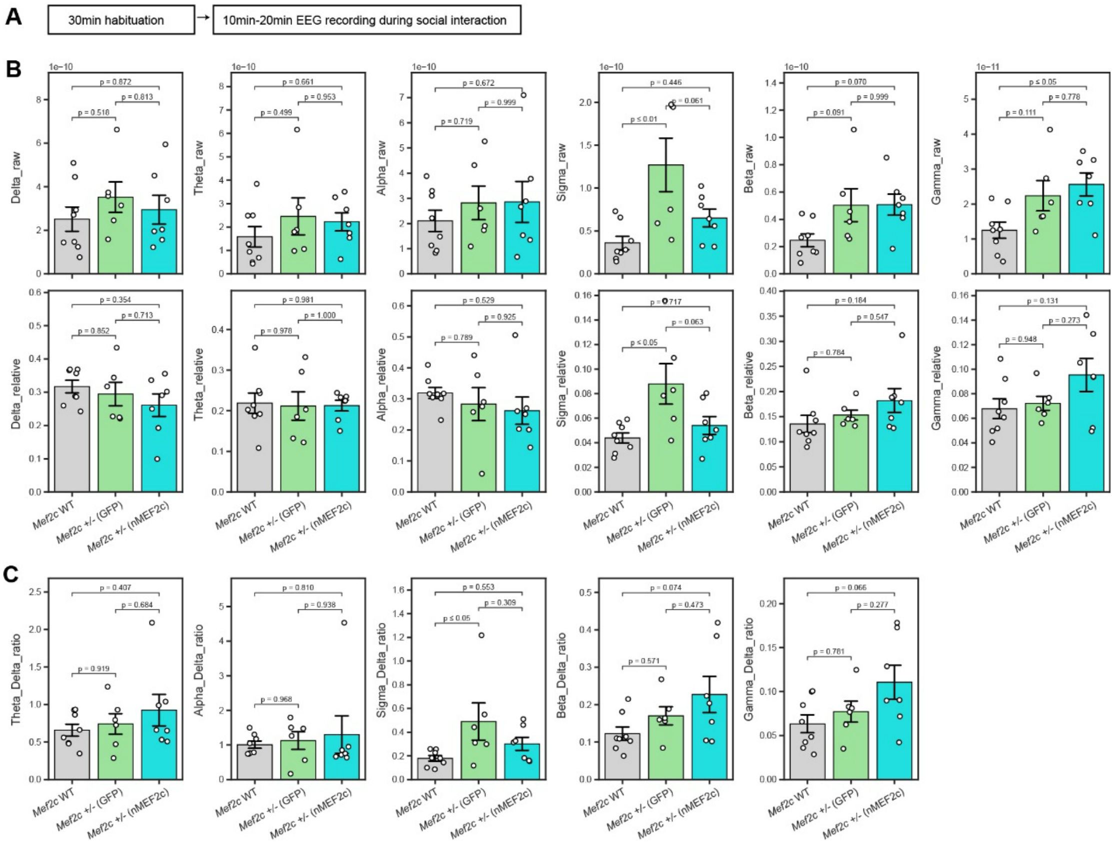


**Supplementary Figure 5. EEG spectral activity during social sniffing following nMEF2C gene replacement.**

(A) Experimental workflow for EEG/EMG recording during social interaction. (B) Absolute and relative EEG band powers during annotated direct social sniffing episodes. (Top) Absolute power in the δ, θ, α, σ (12–15 Hz), β, and γ frequency bands. (Bottom) Corresponding relative power for each band. EGFP-treated *Mef2c*$^{+/-}$ mice exhibited a significant increase in both absolute ($P \leq 0.01$) and relative ($P \leq 0.05$) σ power compared with WT mice. Following nMEF2C treatment, absolute and relative σ powers shifted back toward WT levels, showing no significant difference relative to WT mice. Direct comparisons between nMEF2C-treated mutants and EGFP-treated mutants showed a trend toward reduced absolute ($P=0.061$) and relative ($P=0.063$) σ power that did not reach statistical significance. Absolute γ power was significantly elevated in nMEF2C-treated mice compared with WT mice ($P \leq 0.05$), but did not differ from EGFP-treated mutants ($P=0.778$). No other spectral measures showed significant differences among groups. (C) Power ratios calculated relative to δ activity (θ/δ, α/δ, σ/δ, β/δ, and γ/δ) during social sniffing episodes. EGFP-treated *Mef2c*$^{+/-}$ mice displayed a significantly elevated σ/δ ratio compared with WT controls ($P \leq 0.05$). Treatment with nMEF2C restored the σ/δ ratio toward WT values, rendering the difference from WT mice non-significant. Direct comparison of the σ/δ ratio between nMEF2C-treated and EGFP-treated mutant mice was also non-significant ($P=0.309$). No significant differences were detected across groups for the remaining power ratios (θ/δ, α/δ, β/δ, and γ/δ). Bars show mean ± SEM, and each circle represents one mouse (wild-type, n = 8; EGFP-treated *Mef2c*$^{+/-}$, n = 6; nMEF2C-treated *Mef2c*$^{+/-}$, n = 7). Groups were compared separately for each spectral measure using one-way ANOVA followed by Tukey's multiple-comparisons test. Exact *P* values or significance thresholds are shown.

**Supplementary Figure 6**

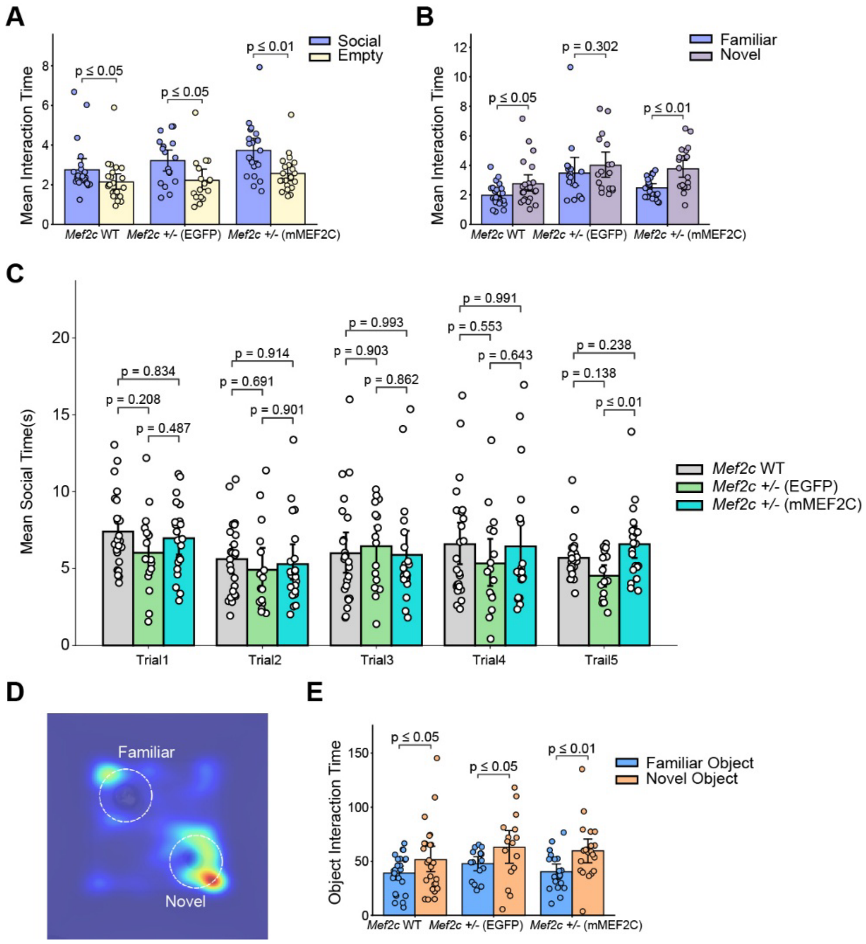


**Supplementary Figure 6. Additional social-interaction analyses and novel-object recognition in the mMEF2C cohort.**

(A) Mean interaction time with the social stimulus versus the empty enclosure during the social-approach phase of the three-chamber test. All three groups showed a significant preference for the social stimulus. (B) Mean interaction time with the familiar versus novel conspecific during the social-novelty phase. Wild-type and mMEF2C-treated *Mef2c*$^{+/-}$ mice showed significant novel-conspecific preference, whereas EGFP-treated *Mef2c*$^{+/-}$ mice did not. (C) Mean social-interaction time during four repeated encounters with the same intruder (trials 1–4), followed by an encounter with a novel intruder (trial 5). During trial 5, mMEF2C-treated *Mef2c*$^{+/−}$ mice showed significantly longer mean social-interaction time than EGFP-treated mutants. No significant between-group differences were detected during trials 1–4. (D)Representative occupancy heatmap during the novel-object recognition test. Dashed circles indicate the familiar and novel object locations. (E) Time spent investigating the familiar and novel objects. All three groups showed a significant preference for the novel object. Groups comprised wild-type mice, EGFP-treated *Mef2c*$^{+/−}$ mice, and mMEF2C-treated *Mef2c*$^{+/−}$ mice. Bars indicate mean ± SEM, and individual points represent individual mice. Brackets indicate statistical comparisons, with exact P values or significance thresholds shown. EGFP, enhanced green fluorescent protein; WT, wild type.
